# Sex-specific non-linear DNA methylation trajectories across aging predict cancer risk and systemic inflammation

**DOI:** 10.1101/2025.08.19.671184

**Authors:** Robin Grolaux, Macsue Jacques, Bernadette Jones-Freeman, Steve Horvath, Andrew Teschendorff, Nir Eynon

## Abstract

Aging is a multi-modal process, leaving distinct signatures across molecular layers, including the epigenome. DNA methylation changes are among the most robust markers of biological aging. Yet, most studies rely on models assuming linear relationships with age and often analyze mixed-sex cohorts, overlooking well-known sex differences in the timing and nature of aging phases. Such approaches risk obscuring critical, non-linear transitions and sex-specific trajectories that may better capture the biology of aging. We developed a computational approach to detect complex, non-linear trajectories and disentangle shared from sex-divergent patterns. Applied to whole-blood deconvoluted methylomes from 252 females and 246 males spanning ages 19–90 years, this analysis revealed convergent and divergent epigenetic aging pathways independent of immune cell composition. These non-linear trajectories were enriched for developmental transcription factor binding motifs, including NF1/CTF and REST, which are known for their oncogenic potential. Strikingly, a female-specific non-linear cluster was robustly associated with cancer onset and systemic inflammation. Our results uncover sex-specific, non-linear aging programs that better capture the dynamics of epigenetic change than linear models. These findings nominate candidate biomarkers for early disease risk and offer mechanistic insight into how aging trajectories diverge between the sexes.

## Introduction

Biological aging is often modeled as a linear process; yet many molecular and physiological changes accelerate, decelerate, or shift phases with age rather than progressing uniformly^1^. This concept of biological non-linearity, changes that deviate from a constant rate over time, is increasingly supported across diverse molecular and physiological domains. Telomere attrition, a hallmark of aging, follows a non-linear trajectory, with faster shortening in early life and slower decline in later decades^2^. Transcriptomics studies in mice have identified cross-tissue non-linear gene expression patterns that correlate with protein expression^3^, as well as late-life shifts in skeletal muscle expression consistent with the ‘elbow’ (i.e. inflection point), typical of non-linear functions^4^. In humans, age-related non-linear patterns have been reported for circulating microRNAs^5^, and undulating changes have been observed in the proteome^6^, transcriptome^7^, and metabolome^1^. Together, these findings suggest that many molecular processes follow complex trajectories across the lifespan, challenging the notion of steady, uniform aging.

DNA methylation (DNAm) is a primary hallmark of aging, and one of the most reliable molecular surrogates for estimating biological age^8,9^. While many epigenetic clocks appear to have a linear relationship with chronological age, their underlying regression models often assume a more complex, log-linear relationship: notably, the Horvath 2013 pan-tissue clock and the Lu 2023 pan-mammalian clock were built by regressing DNA methylation data on a log-transformed version of age, which suggests that methylation changes rapidly in early life and then slows to a constant rate after adulthood^10,11^. While these regression modeling approaches recognize the non-linear nature of methylation aging, they constrain trajectories to simple monotonic forms and cannot capture more complex patterns such as U-shaped curves, multi-phase dynamics, or abrupt inflection points. Indeed, lifespan analyses have documented non-linear DNAm changes at specific CpG sites^12–15^.

Measurements of increased variance, such as variably methylated positions, also exhibit non-linear age-related patterns, highlighting that both the mean and variability of DNAm can shift in complex ways over time^13,16,17^. Despite this, most DNAm studies continue to rely on linear regression or other monotonic models, which preferentially detect features that increase or decrease steadily, while potentially missing, or mischaracterizing, non-linear signals ^18^.

Sex-specific differences in aging are widely recognized at physiological and clinical levels ^19–21^, yet remain underexplored in the context of DNAm. When considered, sex is most often modeled as an interaction term in linear frameworks, limiting the detection of patterns that differ in shape, timing, or magnitude between sexes ^1,14,20,22^. Non-linear approaches are particularly well-suited to uncover such differences, as they can reveal age windows of abrupt divergence, sex-specific inflection points, or distinct multi-phase dynamics.

Here, we investigated sex-specific non-linear aging patterns of DNAm across the adult human lifespan. To enable this, we developed SNITCH (Semi-supervised Non-linear Identification and Trajectory Clustering for High-dimensional data), a robust framework for detecting and clustering CpG sites with shared linear and non-linear age-related changes. We first validated SNITCH using simulated data, then applied it to a high-quality whole-blood DNAm dataset (n=238 males, and 256 females; 18-90 years old), accounting for immune cell composition. This analysis identified both sex-dependent and sex-independent non-linear aging trajectories. Replication in a large independent cohort confirmed that specific non-linear clusters are predictive of inflammation and cancer onset in a sex-specific manner. Our findings demonstrate the value of systematic non-linear analysis for uncovering previously hidden dimensions of epigenetic aging and provide new candidate biomarkers for disease risk stratification.

## Results

### SNITCH: Semi-supervised approach to cluster CpGs based on their aging pattern

Given the widespread evidence of non-linear aging dynamics and the shortcomings of linear analyses, there is a clear need for new methods to detect and characterize non-linear patterns of aging. Understanding aging’s complex trajectory requires analytical approaches that can capture inflection points, accelerations/decelerations, and multiphase changes in biological data. By moving beyond linear models, researchers can unveil previously hidden aging signals. Previous attempts to identify non-linear changes in DNAm have relied on the binning of age categories^14^ or a priori shape of aging patterns^13^. However, systematic tools to scan for arbitrary non-linear trajectories, without pre-specifying a particular model, are needed to truly let the data reveal aging’s patterns. The closest attempt to answer this need has been described by Okada et al.^18^ where functional data analysis was used to cluster CpGs as linear increasing (LI), linear decreasing (LD), non-correlated (NC), or non-linear (NL). Nevertheless, this method remained limited in its ability to discriminate between non-linear trajectories and identify increased variance (VI).

To address this gap, we developed and applied SNITCH (Semi-supervised Non-linear Identification and Trajectory Clustering for High-dimensional data), a heuristic-based statistical framework that distinguishes between linear, non-linear, variable, and non-correlated methylation trajectories. The method leverages both generalized linear modeling and generalized additive modeling to identify CpGs exhibiting distinct age-associated patterns (NC, LI, LD, NL, and VI) while controlling for potential confounders. Unsupervised clustering is then applied on functional principal components of the non-linear positions to highlight CpGs sharing similar non-linear trajectories (Figure 1A). We benchmarked the classification accuracy of SNITCH, in combination with clustering of non-linear patterns, against three stand-alone unsupervised clustering algorithms and the functional trajectory-based DICNAP method^18^. Performance was evaluated using Adjusted Rand Index (ARI) and Adjusted Mutual Information (AMI), using simulated methylation data (i.e., bound by 0-1) from a highly varied pool of distributions with known ground-truth labels (Figure 1B, Methods). We found that SNITCH outperformed stand-alone unsupervised clustering algorithms and DICNAP (Figure 1D & Supp. Fig. 1A). The best results were achieved by using SNITCH + Fuzzy/HDBSCAN (ARI: 0.97; AMI: 0.98). In this setting, we observed a robust concordance between predicted and ground truth labels, with the main missclassifications occurring between the logarithmic decreasing and linear decreasing groups (Figure 1C & Supp. Fig. 1B).

**Figure 1:**
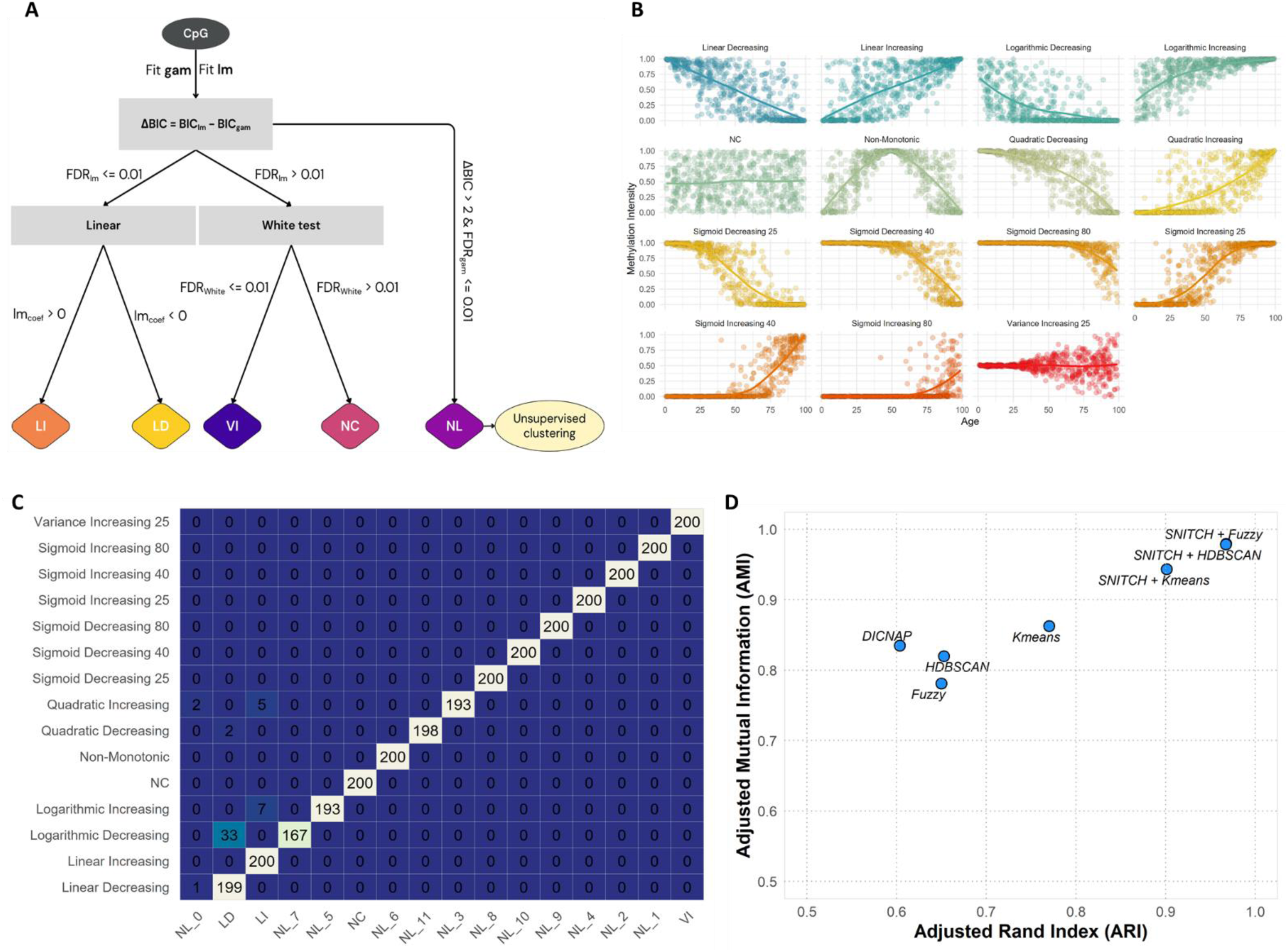
**A**, SNITCH Pipeline. **B**, Distribution of the simulated aging patterns. **C**, Confusion matrix between the ground truth and predicted clusters. **D**, Benchmark of SNITCH compared to stand-alone unsupervised clustering methods and DICNAP. Gam = Generalized Additive Model; lm = Linear Model; BIC = Bayesian Information Criterion; FDR = False Discovery Rate; LI = Linear Increasing; LD = Linear Decreasing; VI = Variance Increasing; NC = Non-Correlated; NL = Non-Linear.

### Sex-specific Non-Linear aging patterns in blood

We then applied SNITCH to a high-quality dataset in blood (GSE246337) containing 238 and 256 males and females, respectively. Blood DNAm is highly influenced by immune cell heterogeneity, and adjustment for cell type composition is essential for the identification of epigenetic modifications independent of the immune profile^23^. Immune cell fractions of whole blood can be effectively estimated by the use of DNAm-based deconvolution methods^24^. Thus, in both females and males, we built three different models of aging trajectories accounting for an increasing number of immune cell types. Our baseline model didn’t include any cell fractions. The 7 cell-model was corrected for B-, NK-, CD4T, and CD8T-cells, Monocytes, Neutrophils, and Eosinophils. Finally, the 12-cell model discriminated between naive and mature B-, CD4T-, and CD8T-cells, and added T-regulatory cells and Basophils. This rigorous approach allowed us to identify DNAm aging patterns occurring independently from changes in immune cell fractions.

Overall, CpG classification was highly stable across models in both sexes: In females, 95.9% (N=534,132) retained identical cluster assignments across all three models. This percentage changed only a little in males (95.6%). This was mainly driven by CpGs classified as Non-Correlated (NC) with age (94% in both males and females). Among the remaining CpGs that changed classification, transitions were most frequently directed toward the NC cluster upon inclusion of immune covariates in both females and males (Figure 2A, Supp. Table 1), highlighting how linear and non-linear trajectories are both confounded by changes in the immune profile. The following analyses describe the results from the 12-cell model if not otherwise stated. In both males and females, the majority of CpGs showed no correlation to age (N_fem = 543,972 (97.7%); N_m = 548,935 (98.6%)). Whereas the number of Non-Linear (NL) CpGs was markedly different between females (N=1305) and males (N=155) (Figure 2B, Supp. Fig 1C). Those results were surprising in light of a recent meta-analysis showing that almost half of the blood CpGs showed differential methylation with age^17^. We investigated whether our analysis was underpowered by combining male and female participants, effectively doubling the size of the cohort, and accounting for sex as a covariate in the model. Consistent with our sex-specific analysis, we found that 93% of the CpGs (N = 516836) stayed non-correlated with age, suggesting that sample size plays a negligible part in our results. These contrasting findings may be explained by our strict QC and significance thresholds (Methods), the high quality of our dataset, and the correction for 12 immune cell fractions against the 5 included in the meta-analysis.

**Figure 2:**
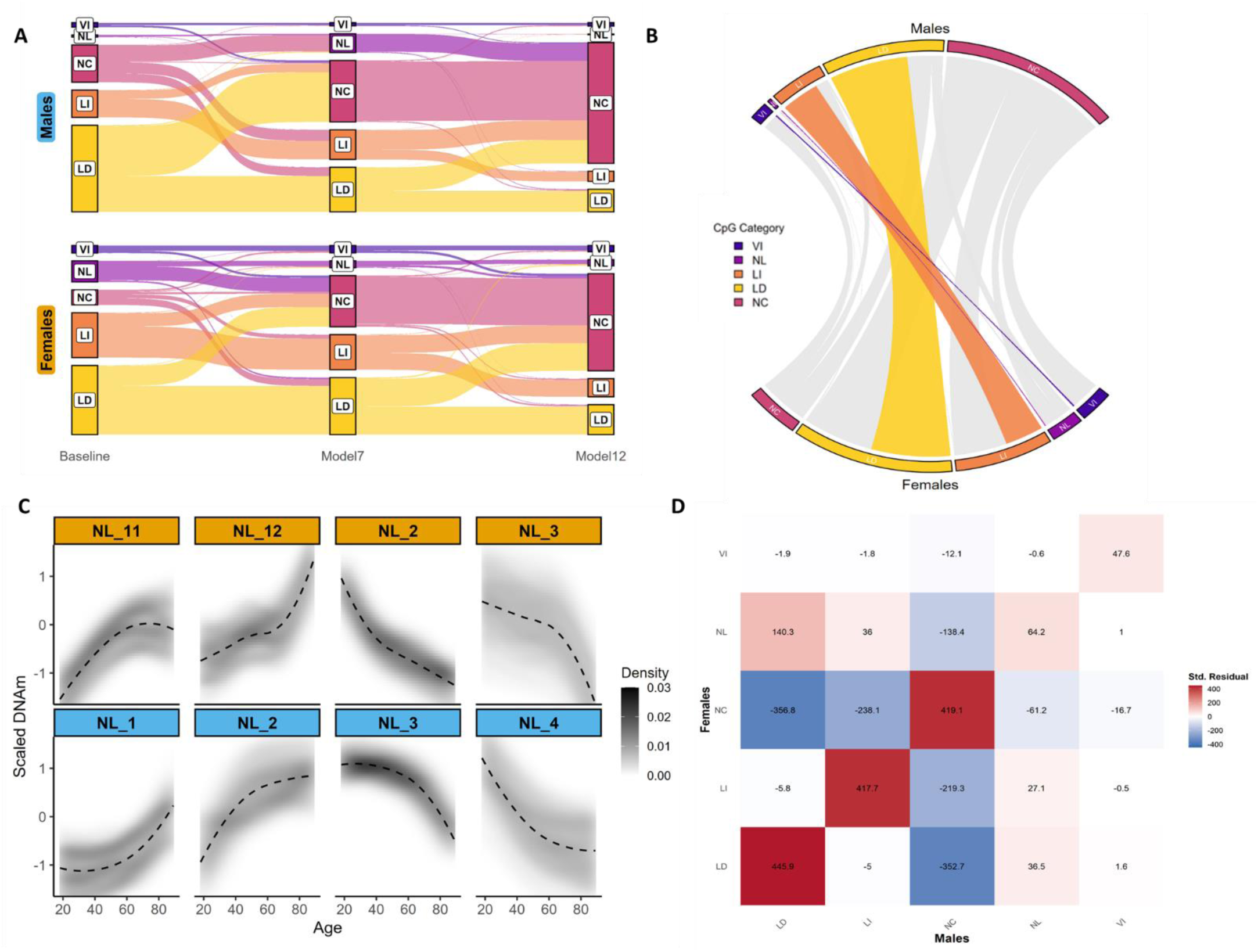
**A**, Repartition of the number of CpGs among clusters in females and males. **B**, Conserved CpGs between male and female clusters. **C**, Non-linear clusters identified in females (top) and males (bottom). Beta values were centered and scaled prior FPCA and unsupervised clustering and are used to better illustrate the patterns. **D**, Chi-square test for the conservation of CpGs among female and male clusters. LI = Linear Increasing; LD = Linear Decreasing; VI = Variance Increasing; NC = Non-Correlated; NL = Non-Linear. Note: In A & B, CpGs that were NC across all 3 models were removed for visualization purposes.

To evaluate the concordance of CpG aging trajectory classifications between sexes, we compared SNITCH-assigned trajectory labels in males and females using a Chi-square test of independence. The analysis revealed a highly significant association (χ² = 403,770, df = 16, pvalue < 2.2 × 10⁻¹⁶), indicating that CpGs were classified into the same trajectory category in both sexes more frequently than expected by chance. Out of the total CpGs assessed, only 9,938 CpGs (∼1.78%) changed trajectory class between males and females, confirming a high degree of overall consistency. This was further supported by the standardized residuals (Figure 2D), which showed strong positive values along the diagonal of the contingency matrix, reflecting substantial overlap in classification across sexes, including in the NL–NL cell, indicating an enrichment of the 39 CpGs classified as non-linear in both sexes despite the discrepancy in the total number of NL CpGs identified. Conversely, the most pronounced negative residuals were observed in off-diagonal cells where CpGs were classified as age-associated (LD or LI) in one sex but NC in the other. This pattern suggests a subset of CpGs with potential sex-specific sensitivity to age-related methylation changes. In addition, NL CpGs showed moderate positive residuals when aligned with LD (+140.3) and LI (+36.0) in the opposite sex, suggesting that a subset of CpGs classified as non-linear in one sex may appear more linear in the other. Finally, the NL–NC cells exhibited a residual of –138.4 in females and −61.2 in males, indicating that CpGs classified as non-linear in one sex were rarely non-correlated in the other, further supporting their functional relevance. To identify clusters of CpGs sharing similar aging trajectories, we applied the last step of our pipeline by performing unsupervised clustering on the functional components of the NL CpGs (Methods). This revealed similar-shaped trajectories in males and females, with 4 primary clusters identified in females and 5 in males (Supp. Fig. 2). To avoid redundancy between similar clusters, we merged clusters with a Spearman correlation coefficient > 0.9 (Supp. Fig. 3). This resulted in a final number of 4 principal NL patterns in females and males (Fig. 2C). Within those, we observed different inflection points, marking a change of pace in the methylation trajectory. In females, clusters 3, 11, and 12 showed an elbow between the 70-80 years mark, where cluster 2 showed it earlier, around the 50 years mark. In males, the inflection points appeared around 60 years for clusters 1 and 3 and 50 years for clusters 2 and 4. This analysis highlighted the similar aging patterns seen in males and females, but hinted at different temporalities for the inflection points.

### Functional analysis of Age-related trajectories

#### Trajectories of clocks’ CpGs

The first step we took to understand the functional role of the clusters we identified was to investigate the classification of CpGs previously used to train epigenetic clocks. A ubiquitous tool in the field of aging, epigenetic clocks are biomarkers that show relevance in assessing the onset of several age-related conditions as well as the utility of therapeutic strategies.

Most epigenetic clocks are machine-learning models based on linear regression^25^ (e.g., elastic-net). By construction, this limits the granularity of the aging trajectories they capture by either overlooking non-linear patterns or over-simplifying them as linear, thus limiting their biological interpretability. Further complicating their interpretation, blood-based epigenetic clocks often ignore cell-type heterogeneity when considering age-related DNAm changes. Within our two cohorts, we looked at the classification labels of the CpGs underlying 9 of the most common clocks^11,26–31^ across our 3 models (Figure 3A). As expected, we found that the proportions of CpGs classified as NC increased between each model iteration, reflecting that part of the signal captured by these clocks arises from age-driven changes in immune-cell fractions^8,25^. In addition, our results highlighted that the majority of the clocks’ CpGs remained classified as NC in our baseline model. Notably, the Hannum clock was the only clock that showed a higher proportion of CpG associated with age (LI & LD) compared to NC across all three models. This finding is consistent with the inherent designs of each clock: the 2nd-generation clocks (PhenoAge, GrimAge, DunedinPace, Zhang_10, and YingAge) were trained on phenotypic age or mortality risk^26,28–32^, overlooking chronological CpGs, and Horvath’s clock was trained across multiple tissues^11^, where Hannum’s clock was built to estimate chronological age in blood^27^. Finally, we observed that most of the clocks captured NL and VI CpGs in males and females. To complement this analysis, we examined a published set of 350 age-associated CpGs shared across immune cell types^33^. Applying the same approach to these loci yielded classifications of hyper- and hypo-methylation that aligned with those reported in the original study, reinforcing the validity of our modeling strategy, which incorporates immune cell composition (Supp. Fig. 4A). Moreover, our results revealed that a subset of these CpGs exhibited non-linear associations with age.

**Figure 3:**
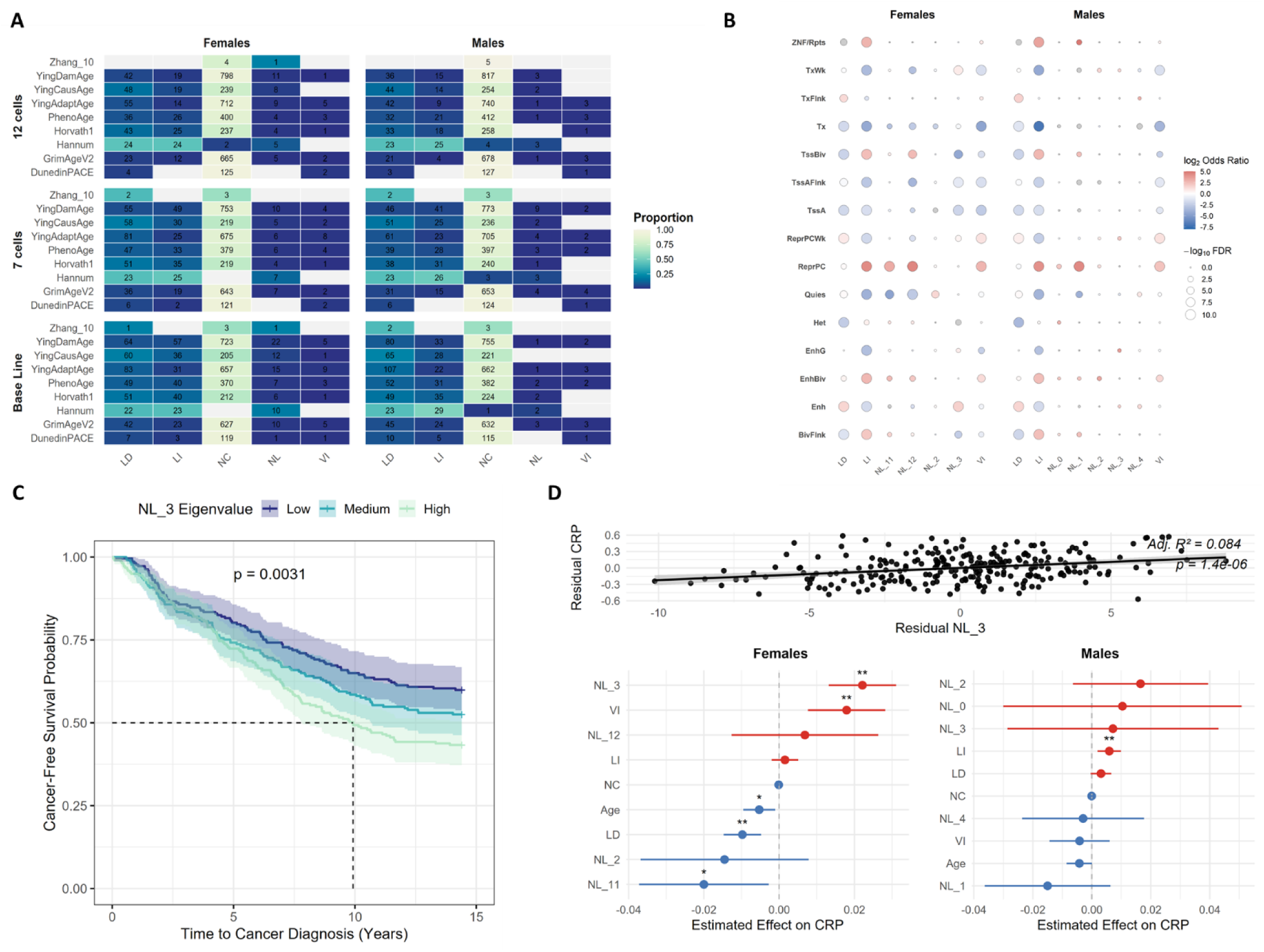
**A**, Enrichment of CpG labels across 9 epigenetic clocks. **B**, Chromatin state enrichment analysis across age-related clusters identified in females and males. **C,** Kaplan–Meier survival curves stratified by NL3 cluster tertiles in females. Participants with high NL3 scores had a significantly shorter cancer-free survival period compared to those in the low and mid tertiles (log-rank p = 0.003). The dashed lines represent the median time before diagnosis. Shaded regions indicate 95% confidence intervals. **D**, Top: Regression model of the female NL3 cluster eigenvalue against estimated CRP levels. Bottom: Coefficient from a linear regression model predicting estimated CRP levels with each cluster’s eigenvalues in males and females. Abbreviations in B correspond to the classes from the Roadmap Epigenomics project^34^. CRP: C-Reactive Protein.

### Functional analyses of sex-specific aging patterns

After successfully identifying sex-specific ageing methylation patterns, we performed separate functional analysis in males and females for all age-associated clusters.

#### Chromatin enrichment analysis

We performed chromatin state enrichment analysis in males and females to determine the epigenetic context of age-associated methylation clusters. Each cluster was tested for enrichment across 15 chromatin states from the Roadmap Epigenomics Project^34^ using the NC cluster as a reference (Methods). Overall enrichment results were highly similar between males and females in the LD, LI, and VI clusters (Figure 3B, Supp. Table 2). Those similarities are consistent with the overlap of CpGs observed in those clusters between males and females (Figure 2C & 2D). Among the remarkable trends, we observed a notable enrichment of the Repressed polycomb and bivalent chromatin states in clusters characterized by increased methylation or age-related variance (LI, VI, NL11, NL12 in females & LI, VI, NL0, NL1 in males). This observation is consistent with previous findings that age-related hypermethylation occurs preferentially in those domains^35–37^. Overall, the enrichment of chromatin states at NL clusters seemed mostly driven by the directionality of the trajectories (gain vs. loss of methylation) rather than by the individual patterns, as they were concordant with the LI or LD clusters, respectively. A noticeable deviation from this was the significant enrichment of “weak transcription” states in NL3 in females (hypomethylated with age) compared to its depletion in LD (NL3 log₂ OR = 0.7, LD log₂ OR = −0.14). This is supported by previous studies highlighting that hypomethylated sites are enriched in active or transcribed genomic regions, including the weak transcription chromatin state^38,39^. In particular, this suggests a breaking point around 60 years old in females, where erosion of DNA methylation in weakly transcribed or formerly silent chromatin regions could lead to leaky transcription, supporting the theory of increased transcriptional noise in aging^40^.

#### Pathway enrichment analysis

Next, we performed pathway enrichment analysis for our non-linear clusters using different databases (the Gene Ontology (GO) Molecular Function (MF) and Biological Process (BP), Kyoto Encyclopedia of Genes and Genomes (KEGG), and Reactome). Enrichment analyses in DNA methylation datasets are inherently biased toward long genes, and due to CpGs mapping to multiple genes^41,42^. To account for those, we used *missmethyl,* a method that addresses these biases by leveraging prior probabilities^42,43^. Across all NL clusters, only NL12 in females was enriched for the term “Neuroactive ligand signaling” in KEGG (FDR = 0.023); no enrichment was observed for NL clusters in males (Supp. Table 3).

#### Motif enrichment analysis

In addition to the local chromatin context, the underlying DNA sequence also contains biologically relevant information that can help understand the function of a particular set of CpGs. Although the mechanisms haven’t been fully elucidated yet, it is widely accepted that the methylation context at a specific motif can positively or negatively impact binding affinity of transcription factors (TFs)^44,45^. Focusing on the NL CpGs, we performed cluster-wise motif enrichment analysis to identify the presence of TFs binding sites (TFBs) in their vicinity (Methods). In females, only NL12 and NL3 showed significant enriched motifs (Supp. Fig. 4B, Supp Table 3). NL12 was enriched in ZNF652 binding site (qval = 0.0249) where NL3 was enriched for both Nuclear Factor 1 (NF1/CTF) half-(qval = 1×10^-5^) and full motif (qval = 0.0026), Hoxc9 (qval=0.0411) and Gata6 (qval = 0.0476). In males, only NL1 and NL4 were enriched in TFBs (Supp. Fig. 4C, Supp. Table 4). Similar to NL3 in females, the NL4 cluster in males was enriched for NF1 half-(qval = 0.0063) and full-site (qval = 1×10^-4^) along with REST/NRSF (qval = 1×10^-5^). The NF1-CTF family of transcription factors was shown to regulate cell development in the central nervous system, is associated with cancer, and plays a key role in the regulation of transcription^46–48^. Similarly, ZNF652 acts as a potent tumor suppressor in breast cancer by repressing the transcription of oncogenes^49,50^, while both Gata6 and Hoxc9 are involved in embryogenesis and have oncogenic properties^51–54^.

### Nonlinear clusters as biomarkers of diseases

#### Cancer risk

DNA methylation, or surrogate measures such as epigenetic age-acceleration, have been used as biomarkers to predict the onset of or diagnose a wide range of pathological conditions, ranging from rare developmental diseases to cancer^55–58^. Given the presence of transcription factor binding motifs known to regulate oncogenic pathways within the methylation clusters, we hypothesized that these epigenetic modules may capture pre-diagnostic signals of cancer risk. To evaluate the predictive value of the NL clusters for cancer onset, we assessed the association between their eigenvalues and cancer development using three complementary approaches: Cox proportional hazards models, Kaplan–Meier survival analysis, and logistic regression. Analyses were conducted in the EPIC-Italy cohort^59^, a large prospective study in which blood DNA methylation was profiled at baseline in healthy participants, with up to 15 years of follow-up for incident cancer diagnoses. The cohort includes time-to-diagnosis information for several cancer types, with breast cancer (C50) and colorectal cancer (C18) being the most prevalent. We first assessed associations across all cancer types. Stratifying by sex revealed a clear sex-specific predictive capacity. In females, NL3 emerged as the most robust predictor of cancer risk across all analyses. In Cox regression, NL3 eigenvalues were significantly associated with a shorter time to cancer diagnosis (HR 1.020, 95% CI: 1.007–1.032, FDR = 0.0058) (Supp. Table 4). This was supported by Kaplan–Meier survival analysis, where NL3 tertiles showed clear separation in cancer-free survival (log-rank p = 0.003, Figure 3C). Logistic regression confirmed the predictive effect (OR = 1.032, 95% CI: 1.012–1.052, FDR = 0.0083). The NL11 cluster also showed a potential association with cancer risk. While its Cox result was borderline (HR = 1.039, FDR = 0.063), logistic regression yielded a significant effect (OR = 1.065, 95% CI: 1.008–1.126, FDR = 0.049), suggesting a possible predictive role that warrants further investigation. No associations were detected for NL2 or NL12 in any models. In males, none of the tested clusters showed significant associations. Next, we stratified the cohort by cancer type to determine whether these associations were driven by specific tumor types. In the breast cancer sub-cohort (female participants only), NL3 remained a robust predictor of disease onset in both models (Cox HR = 1.023, 95% CI: 1.009–1.037, FDR = 0.0045; Logistic OR = 1.035, 95% CI: 1.013–1.058, FDR = 0.0051) (Supp. Table 4). Likewise, NL11 showed consistent predictive value (Cox HR = 1.065, 95% CI: 1.019–1.113, FDR = 0.0097; Logistic OR = 1.101, 95% CI: 1.034–1.172, FDR = 0.0051). These results suggest that NL3 and NL11 capture biologically relevant, pre-diagnostic methylation signals specifically associated with breast cancer development. In contrast, no non-linear cluster was significantly associated with colorectal cancer (C18) in either males or females. Both Cox and logistic regression models yielded non-significant associations across all clusters, indicating that the epigenetic trajectories captured by these modules do not predict colorectal cancer onset within this cohort. Together, these findings highlight the sex- and cancer-type-specificity of non-linear DNA methylation aging patterns. While NL3 and NL11 showed significant biomarker potential for breast cancer risk in females, their lack of association with colorectal cancer suggests these epigenetic trajectories reflect tissue-specific aging dynamics rather than generalized cancer susceptibility.

#### Inflammation - CRP levels

Senescence of the immune system is termed “inflammaging”, with chronic inflammation being one of the hallmarks of aging^60,61^. Particularly relevant to our blood-based analysis, we decided to examine the association between the non-linear clusters and inflammation. We explored whether some of the age-related clusters we identified shared this predictive capacity. First, we looked at inflammation. Separately in males and females, we calculated each cluster’s eigenvalues and looked at their association with estimated C-reactive protein (CRP) levels (Methods), a well-established marker of inflammation^62^. Three nested models were used to assess these associations while adjusting for age and NC CpGs (Methods). The analysis revealed sex-specific patterns of association, with several eigenvalues significantly predicting CRP independently of chronological age. In females, the full linear model including eigenvalues from all age-related clusters significantly improved CRP prediction compared to age + NC alone (*adjusted R² =*0.422; ANOVA *p* < 8.2 × 10^-11^). Several modules showed robust associations (Figure 3D, Supp. Table 4). The NL3 module displayed the strongest positive association with CRP (β = 0.0221, *p* = 2.06 × 10⁻⁶), suggesting that methylation patterns in this cluster closely align with inflammatory status. The LD module was significantly negatively associated with CRP (β = –0.0098, *p* = 1.34 × 10⁻⁴), indicating a potential protective or anti-inflammatory methylation pattern. NL11 also revealed a significant negative association (β = –0.0200, *p* = 0.023), while VI showed a significant positive association (β = 0.0179, *p* = 6.9 × 10⁻⁴). Notably, the NC eigenvalue was not significantly associated with CRP (*p* = 0.266), although it contained 87% of the CpGs used in CRP estimations (Supp. Fig. 4D). This result likely stems from the fact that we accounted for 12 immune cells fractions in our model, including naïve and mature Tells which the CRP signature relies on^62^, and further shows that the NL patterns captured methylation variation independently of the immune profile. Together, these results suggest that inflammation in females is selectively captured by distinct age-related methylation patterns, some of which are positively associated with inflammatory burden (e.g., VI, NL3) and others inversely associated (e.g., LD, NL11), reflecting complex and potentially compensatory epigenetic dynamics. In contrast, males exhibited a more restricted profile of significant associations. While the full model remained statistically significant (*adjusted R²* = 0.423; ANOVA *p* = 4.82 × 10^-^⁵), only the LI module showed a significant positive association with estimated CRP levels (β = 0.0059, *p* = 0.0037) (Figure 3D, Supp. Table 4). Associations with other clusters, such as NL1, NL2, and LD, did not reach significance, although some trends were observed. The NC module was again not predictive (*p* = 0.65), reinforcing the specificity of the signal to age-related modules. These sex-specific patterns suggest that while methylation-based inflammation signatures exist in both sexes, females display a more modular and interpretable architecture. In contrast, the inflammatory signal in males may be either more diffuse across the methylome or less tightly coupled to the cluster structure identified in the present analysis. Together, these findings indicate that methylation clusters derived from non-linear age-related trajectories carry relevant information about the inflammatory burden in blood.

### Different waves of regulation in males and females

In the analysis above, we identified functional clusters of age-related CpGs sharing similar linear or non-linear patterns. Although this analysis hinted at specific ages at which disruption of methylation trajectories occurs (i.e., inflection points), another analytical method has been previously used to address this question directly^1,6^. We applied the same method used by Shen & al.^1^ to reveal peaks of disruption of methylation patterns in our male and female populations (Methods). Our analysis focused on age-related CpGs identified at the last step and revealed distinct waves of dysregulation in males and females (Fig. 4A). Those peaks were deemed robust as they passed different significance thresholds (Fig. 4B). In females, we observed three peaks at age 33, 51, and 73 that were consistent with 2 of the inflection points observed in the previous analysis. Whereas males only displayed two peaks at 47 and 63 years old, again consistent with the clustering analysis.

**Figure 4:**
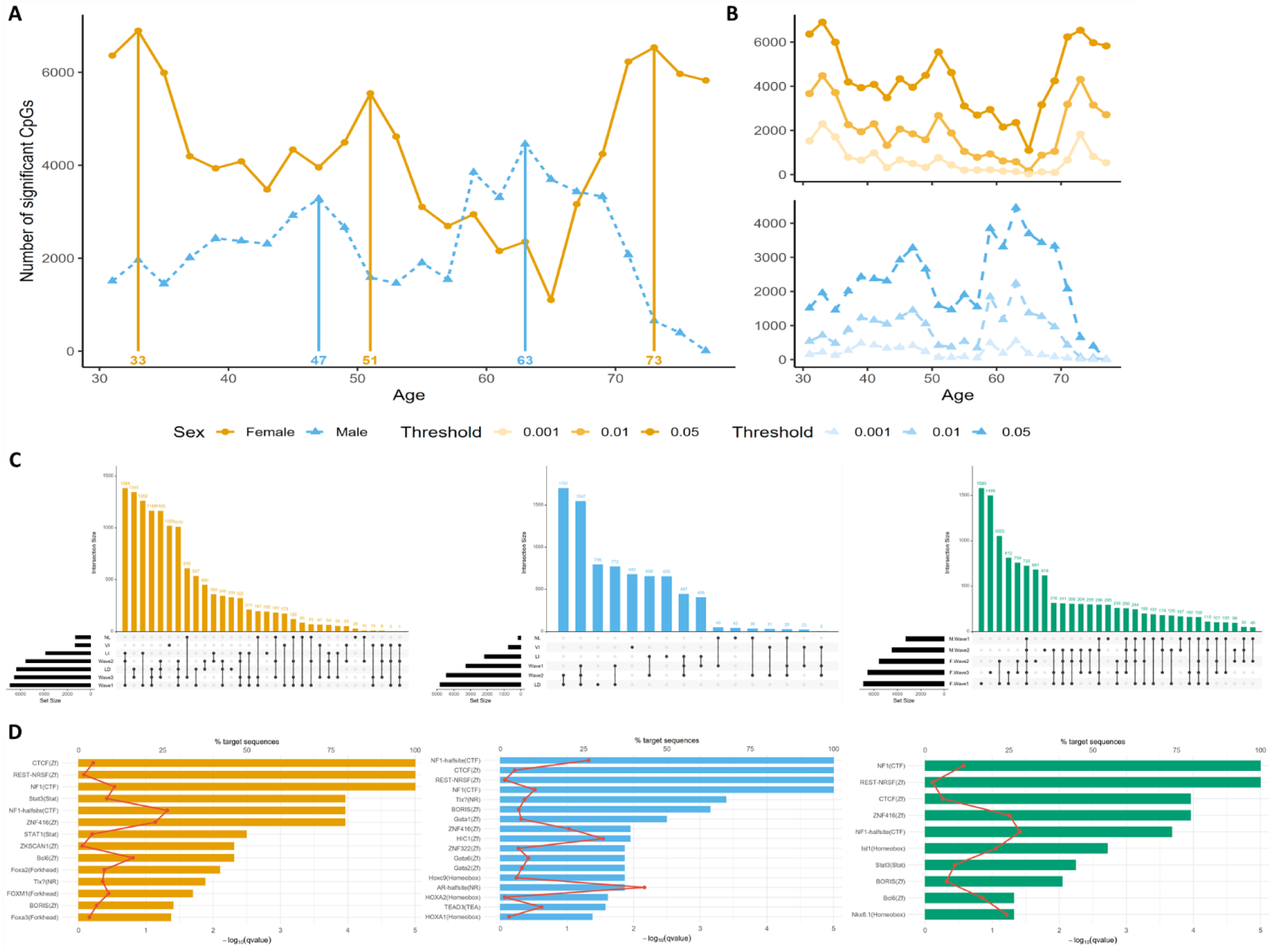
**A**, DEswan analysis in males and females. Each point represents the number of CpGs dysregulated between a 15-year windows on each side of the specified age at FDR < 0.05. **B**, DEswan analysis at different FDR thresholds. **C**, The overlapping CpGs between waves of regulations and NL clusters in females (left), males (center), and between male and female waves (right). **D**, Motif enrichment analysis among common CpGs identified across waves of dysregulation in females (left), males (center), and across sex (right).

The number of significantly dysregulated CpGs markedly differed between males and females, with females showing an overall higher number, similar to the discrepancy observed between their respective numbers of NL CpGs. Notably, the peaks at 33 and 51 in women are supported by previous findings highlighting those decades as transitional periods in women’s aging^63^ and previous research in protein expression levels identified three peaks at 34, 60, and 78 years old^6^. Recent findings in a multi-omics setting also identified crests of dysregulation around 40 and 60 years old. Both those analyses were performed in a joint cohort of men and women and might represent a combination of the sex-specific peaks we identified. Thus, our results support previous claims that aging occurs in waves across multiple modalities rather than unique molecular components. In addition, we highlighted a distinct temporality in the aging process across sex, as underlined by the earlier onset of the crests in males. Integrating this analysis with the clusters we identified previously revealed that the main CpG overlaps in males and females occur with the linear clusters LI and LD (Figure 4C). Most of the NL CpGs in females intersected with wave 3, whereas in males, they were equally represented in waves 1 and 2. This analysis further highlighted conserved CpGs across waves (N_males = 2027 & N_females = 1453) and across sexes (N = 725). Motif enrichment analysis of those sets confirmed the enrichment of NF1 full- and half-binding motifs at age-related CpGs, and revealed the presence of CTCF and NRSF/REST binding sites, two consistent marks of epigenetic aging^10,33,64–67^, among other TFBs (Figure 4D). To understand the specific pathways that could be affected, we repeated our previous enrichment analysis, focusing on each peak’s unique set of CpGs (Methods). In females, waves 1, 2, and 3 showed, respectively, 2057, 873, and 2077 uniquely dysregulated CpGs. Wave 1 showed striking enrichment for neurodevelopmental processes, including *nervous system development* (FDR = 3.9 × 10^-4^), *neuron differentiation* (FDR = 8.5 × 10^-4^), and *generation of neurons* (FDR = 1.8 × 10^-3^) (Supp. Fig. 5). Additionally, terms related to transcriptional control and chromatin binding, such as *DNA-binding transcription factor activity* (FDR = 5.3 × 10^-4^), and *double-stranded DNA binding* (FDR = 4.2 × 10^-3^) were significantly enriched. Wave 2 yielded a single enriched term: *regulation of execution phase of apoptosis* (FDR = 0.036) (Supp. Table). These findings recapitulate previous observations of neurodevelopmental and transcriptional gene dysregulation with age^10,68^. Interestingly, in males, among the unique CpGs in wave 1 (N= 1250) and wave 2 (N=2426), no significant enrichment was found. Likewise, the 725 CpGs common to both sexes showed no detectable enrichment. This highlights sex-specific epigenetic changes in females, with broader and more functionally coherent methylation changes. Notably, REST, a transcriptional repressor enriched in our motif analysis, is known to silence neuronal genes in non-neuronal lineages and is a key regulator of the epigenetic program that maintains cellular identity^67,69^. Its involvement, alongside the dysregulation of neurodevelopmental pathways, supports a model in which age-related methylation drift compromises the fidelity of cell-type–specific gene regulation, allowing partial re-expression of lineage-inappropriate developmental programs. This aligns with emerging evidence from aging transcriptomes and methylomes indicating that immune cells in older individuals exhibit loss of identity and ectopic activation of developmental gene networks, including those tied to nervous system formation^70,71^.

## Discussion

By leveraging a novel analysis pipeline and a high-quality DNA methylation (DNAm) dataset, we identified sex-specific non-linear aging trajectories in blood. Fine-tuning our clustering pipeline on simulated data enabled us to establish robust heuristics facilitating the identification of diverging linear and non-linear patterns (Figures 1A & 1B). We showed that our new pipeline, SNITCH, outperforms stand-alone unsupervised clustering methods in discriminating between Variance Increase (VI), Linear Increasing and Decreasing (LI & LD), Non-Correlated (NC), and various Non-Linear (NL) functions (Figures 1C & 1D). Nevertheless, we observed misclassification between the linear and logarithmic patterns due to their close resemblance. By providing SNITCH as a user-friendly framework, our approach is well-positioned to uncover non-linear, tissue- or context-specific aging dynamics in large cross-sectional or longitudinal datasets, including EWAS, transcriptomic time series, or multi-omic aging studies.

The immune profile reshapes with age, and analyses in whole blood are particularly sensitive to those changes^72^. Applying our pipeline to a blood DNA methylation dataset while accounting for an increasing number of immune cell types highlighted the confounding effect of the immune profile on age-related DNAm changes (Figure 2A).

Stratifying our analyses across sexes revealed a notably higher number of NL CpGs identified in Females (Figure 2B). Nevertheless, CpGs generally showed concordant aging trajectories across sexes, with significant enrichment for matching classifications (Figure 2D). A surprising result was the small number of age-related CpGs identified (< 4%), seemingly contradicting findings of a meta-analysis of age-related DNAm in blood^17^. We argue that our results are a conservative but robust representation of true changes occurring during aging. We used a gold-standard reference DNAm dataset that minimises batch effects, covers an equivalent number of males and females, and has a homogeneous age distribution^73^. This, combined with our strict quality control and FDR threshold and the inclusion of 12 immune cell types as covariates in our model, ensured the identification of robust changes and explains those numbers. However, we acknowledge that due to our limited sample size, we might have missed smaller effects or population-specific changes.

Our study helped characterise CpGs underlying epigenetic clocks. We assessed the distribution of clock CpGs within our different aging clusters and found that most models relied on CpGs classified as NC in our model (Figure 3A). Although counterintuitive, we focused on changes independent of the immune profile, where clocks are susceptible to cell fraction changes. This suggests that many clocks rely on CpGs that are stable across age when accounting for immune composition, potentially reflecting their role as proxies for cellular fraction rather than methylation-only age-related change. This was confirmed by our baseline model (not accounting for immune cells), which showed fewer CpGs classified as non-correlated. Additional explanation comes from the data used to train the clocks.

Indeed, among the 9 models we investigated, only the Hannum clock was trained solely on blood DNAm using chronological age as the target variable, where the other clocks were trained on multiple tissues (Horvath’s) or phenotypic variables (PhenoAge, GrimAge, DunedinPace, Zhang_10, and YingAge). We found that most of the clocks included VI and NL CpGs. Further analyses should investigate the weight associated with those in each model to understand the dependency of the clocks on non-linear trajectories, and potentially fine-tune them to reflect the non-linear phases of aging.

Our analysis confirmed the enrichment of Polycomb Repressed chromatin states at hypermethylated CpGs^35–37^. Overall chromatin state enrichments were dependent on the gain or loss of methylation, irrespective of sex or linearity (Figure 4B). Only the NL3 cluster in females differed from this trend and showed a small enrichment for the *weak transcription* state. Consistent with other results, the motif enrichment analysis revealed an enrichment in the motif of REST-NRSF^66^ in the male hypermethylated NL1 cluster. The loss of REST is associated with cognitive impairment and Alzheimer’s disease^67^, and hypermethylation at its binding sites could disrupt its function. The relevance of this finding in blood should be investigated, as is the biomarker potential of the males’ NL1 cluster for Alzheimer’s disease. This analysis also identified new enriched motifs at NL CpGs for the NF1-CTF family, both in males and females, suggesting a potential disruption of their function. Additional transcription factors (TFs) having their binding sites enriched included the STAT family in males (STAT3 & STAT4), underlying a potential disruption of the immune system ^74^ and developmental TFs in females (HOXC9, GATA6)^51,52^. Notably, the enrichment of motifs from developmental TFs at hypomethylated sites could indicate the loss of a safeguarding mechanism consistent with the epigenetic drift view. Nevertheless, our analyses suggest that this drift is not constant but suffers from episodes of dysregulation.

The functional role of our NL clusters was further explored by looking at their relationship with inflammation and predictive capabilities for cancer onset in an independent cohort. With a robust statistical model accounting for immune cell fractions, we identified NL3 in females as being strongly associated with both inflammation (Figure 3D) and cancer development probabilities (Figure 3C), and the female NL11 cluster as mildly associated. The functional specificity of those clusters was underscored by the absence of predictive power from the NC module across sexes. Those results showed the relevance of our identified NL clusters by validating their functionality in an independent cohort. Furthermore, the more robust and multifaceted associations in females strongly support the argument for sex-stratified biomarkers, particularly in the context of aging, cancer, and inflammation. Our motif enrichment analysis laid the ground for further investigating the mechanisms underlying the predictive capacity of females NL3. Notably, NF1/CTF, Hoxc9, and Gata6 binding motifs were overrepresented in this cluster. Gata6 was recently shown to be part of a central mechanism promoting cancer-associated fibroblasts in breast cancer^54^. The study found that the expression of Gata6 was enhanced by the action of TET1, a protein that removes methylation. Evidence shows that binding of members of the Gata family is disrupted by methylation marks^75^. Thus, the hypomethylation observed in NL3 could be associated with increased Gata6 oncogenic activity. Both Hoxc9 and members of the NF1 family have also been associated with breast cancer, although the mechanisms have not been fully elucidated and are not well documented^53,76,77^. Nevertheless, how the occurrence of these epigenetic marks in immune cells may affect cancer onset remains to be investigated.

Overall, our clustering analysis revealed sex-specific functional clusters of NL CpGs. They hinted at sex-specific peaks of dysregulation by showing inflection points around the 75 (NL3, NL11, NL12) and 50 (NL2) years mark in females against the 50 (NL2, NL4) and 60 (NL1, NL30) years mark in males. We used the previously described DEswan analysis^1^ to formally identify peaks of dysregulation in a sex-specific manner. The DEswan results reinforced the findings of our first analysis by revealing peaks of dysregulation at 51 and 73 years old in females, and 47 and 63 years old in males (Figures 4A & 4B), plus an earlier peak at 33yo in females. Those results are strongly supported by the literature, with studies in proteomics and multi-omics identifying peaks at similar ages^1,6,63^. The sex-specificity of our findings indicates an earlier onset of dysregulation in males, with the last wave occurring a decade before the one in females. This also suggests a peak of dysregulation in mid-adulthood occurring in females but not males, although this last finding should be tempered by the minimum age of 25 in our cohort, probably missing prior peaks during adolescence^78^. In both males and females, CpGs identified at those peaks showed a partial overlap with the clusters (Figure 4C). In addition, both males and females showed CpGs consistently dysregulated at all peaks, with some of those conserved across sexes. Motifs enrichment for sex-specific and non-sex-specific conserved CpGs supported our previous findings with NF1/CTF and REST amongst the most enriched binding sites. Consistent with other findings^65,66^ but absent from our cluster analysis, the CTCF motif was also enriched. Our functional analysis for the waves of dysregulation pointed toward a larger and more coherent dysregulation in females, with previously reported pathways related to DNA binding and neurogenesis being affected.

Our study presents several limitations. First, the datasets lacked critical covariates such as BMI or smoking status, preventing us from fully disentangling lifestyle effects from intrinsic aging-related methylation changes. Additionally, both datasets were predominantly composed of individuals of European ancestry, which may limit the generalizability of our findings to more diverse populations. Despite covering a wide age range (25-90 years old), we missed samples in the early infancy/adolescent stages, known to be associated with wide changes in DNAm^78–80^. As a result, our characterization of methylation dynamics during early development remains incomplete, potentially missing early-life inflection points critical to the trajectory of aging. Another limitation is the cross-sectional nature of our study. This is a typical limitation in the field due to the lack of large longitudinal datasets. Thus, our analysis relies on the hypothesis that DNAm changes are conserved to a level across different individuals, a hypothesis supported by the large body of literature on DNAm in aging. Nevertheless, adopting a longitudinal approach is an important step towards personalised medicine and is a direction the field should embrace. In this work, we chose to work with a smaller, high-quality dataset to reduce noise and uncover true age-related changes. While this strategy facilitated the detection of robust changes validated in an independent cohort, our relatively small sample size (∼240 individuals per sex) may have limited our ability to detect subtle but biologically relevant effects, reflecting the small number of age-related CpGs we found. Finally, DNAm is intrinsically difficult to functionally link to changes in the phenotype, in part due to the variety of effects that gain or loss of methylation can have depending on their location. Consequently, we emphasize that our findings are correlative and should be interpreted as surrogate markers rather than direct drivers of phenotypic changes.

In the future, we aim to further explore non-linear patterns in other tissues^81^ and extend our methodology to other omics, such as transcriptomics and proteomics, where the increase of variance is often omitted from clustering analyses. Our results, along with prior work, reveal enrichment of developmental transcription factor motifs (e.g., REST, NF1/CTF) at age-dysregulated CpGs^36,45^. This observation remains underexplored, particularly regarding how such TF-DNAm interactions shift with age and impact transcriptional regulation. Future studies integrating these findings with transcriptomic or chromatin accessibility data could elucidate the functional consequences of these methylation changes. Thus, we will focus on the female NL3 cluster that showed biomarker potential, working to characterise and validate it in other cancer cohorts.

Altogether, our findings support a paradigm shift in the way we conceptualize aging: not as a steady, linear decline, but as a non-linear and entropic process characterized by temporally confined windows of epigenetic instability. By uncovering sex-specific, wave-like patterns of DNA methylation changes that mirror inflection points reported in other omics layers, our study strengthens the case for integrated, temporal frameworks of aging biology. The critical decades we identified, particularly around the 30s, 50s, and 70s, may represent biologically vulnerable windows, where intervention could yield the most impact. Beyond these insights, our findings underscore the need to move beyond one-size-fits-all models and adopt analytical strategies that reflect the inherent heterogeneity, non-linearity, and sex-specific nature of biological aging. However, to fully harness the translational potential of these observations, a pressing need for molecular validation remains. Future work should prioritize characterising the regulatory mechanisms underpinning these methylation dynamics, including their interaction with chromatin architecture, transcription factor networks, and other epigenetic layers, to distinguish causality from correlation and to illuminate actionable pathways of aging and disease.

## Methods

### Cohort and DNA methylation preprocessing

Two separate cohorts were used in this study. GSE246337^73^ was used in the identification of ageing methylation patterns, and GSE51032 was used to identify biomarkers of cancer. For both datasets, the Raw IDAT files from the Illumina Infinium HumanMethylationEPIC v2 BeadChip array (EPICv2) or 450K, respectively, were retrieved from GEO and processed using the *sesame* R/Bioconductor package (v1.18.1)^82^. To harmonize probe identifiers and accommodate the EPICv2 platform structure, prefix collapsing was enabled during data import. Initial preprocessing followed the “QCDPB” pipeline within *sesame*, incorporating: (i) probe quality filtering using the pOOBAH method to remove probes with poor detection p-values, (ii) dye-bias correction to normalize type I and II probe discrepancies, (iii) masking of probes known to be problematic due to non-specific hybridization or SNP interference as defined by Zhou et al.^83^, and (iv) exclusion of probes supported by fewer than four beads.

The resulting beta value matrix was further filtered to improve data integrity. Probes with missing values in more than 1% of samples were discarded. Subsequently, samples with more than 1% missing beta values across retained CpGs were also removed, and we removed probes on the sex chromosomes. The remaining missing values were imputed using k-nearest neighbors using the impute.knn functions from the *impute* package, keeping the default parameters. Beta values were rounded to three decimals for downstream analyses. For the GSE246337 dataset, we only kept probes present on the EPICv1 array, as most tools used in the downstream analysis are not yet compatible with the EPICv2 array.

The final number of probes used in the downstream analyses was 556811, and final numbers of samples were 256 females and 238 males for GSE246337. For the GSE51032 cohort, the final number of probes was 382717, for 651 females and 186 males.

### Description of the SNITCH analysis pipeline

The SNITCH pipeline involved three main steps: Heuristic-based classification, Functional Principal Components Analysis (FPCA), and unsupervised clustering.

### Heuristic-based classification

For each CpG site, methylation beta values are modeled as a function of chronological age. The core procedure included the following steps:

1. **Model Construction**: Linear models (LMs) are fitted using ordinary least squares regression with age as a continuous predictor (lm function from base R). Parallelly, generalized additive models (GAMs) are fitted using restricted maximum likelihood (REML) and thin plate regression splines (*s(Age, k=5)*), enabling the detection of non-linear trends (gam function - *mgcv*^84^). When relevant, covariates are included consistently across both models to preserve interpretability and comparability.
2. **Model Comparison and Heteroscedasticity Testing**: The explanatory performance of GAMs relative to LMs is evaluated using the Bayesian Information Criterion (BIC) (BIC function from base R). CpGs with ΔBIC (BIC(LM) − BIC(GAM)) > 2 are considered to favor the non-linear model. To characterize potential violations of homoscedasticity that may underlie complex aging dynamics (VMP), the White’s test is applied to each LM, with heteroscedastic CpGs flagged based on a 1% FDR-adjusted significance threshold.
3. **Prediction and Effect Size Estimation**: For CpGs favoring a GAM fit, DNAm values are predicted across a continuous age grid ranging from the minimum age of the cohort to the maximum with a one-year step increase, holding covariates constant at reference levels (medians for numeric or first level for categorical variables). This effectively smoothed the trajectories for subsequent steps. For linear trajectories, the direction and significance of age-associated change are inferred from the LM coefficient and p-value, respectively.
4. **Multiple Testing and Classification**: P-values from LM, GAM, and heteroscedasticity tests are corrected using the Benjamini-Hochberg method. CpGs are classified as:

○ **LI** (Linear Increase): Significant linear association (adj. p(LM) ≤ 0.01) with a positive slope.
○ **LD** (Linear Decrease): Significant linear association with a negative slope.
○ **NL** (Non-linear): ΔBIC > 2 and adj. p(GAM) ≤ 0.01.
○ **VI** (Variance-Increasing): No linear association (adj. p(LM) > 0.01) but significant heteroscedasticity (adj. p(White) ≤ 0.01).
○ **NC** (Non-correlated): CpGs not meeting any of the above criteria.

This classification strategy allows SNITCH to robustly detect a spectrum of epigenetic aging signatures, from canonical linear changes to more complex non-linear patterns. The SNITCH method is available and can be installed as a user-friendly R package (https://github.com/fishrscale/SNITCH), thus facilitating access to non-linear trajectory analyses.

### Functional Principal Component Analysis (FPCA)

To further dissect heterogeneity within non-linear (NL) CpG methylation trajectories, we implemented a two-stage dimensionality reduction and clustering procedure based on functional principal component analysis (FPCA) followed by density-based unsupervised classification. This enabled the grouping of NL CpGs into discrete functional subclusters based on the shape of their smoothed age-associated methylation trajectories.

For all CpG sites previously classified as NL by SNITCH, we use the smoothed beta values predicted by the gam model. FPCA is then applied using the fpca.face() function from the *refund* R package, with age as the functional domain. The number of knots is set dynamically based on the number of time points (min(35, floor(0.8 * timepoints))), and the proportion of variance explained (PVE) threshold is conservatively fixed at 99.99% to retain fine-grained trajectory information. This process decomposes the complex, high-dimensional nonlinear patterns into a reduced set of orthogonal functional basis scores.

### Unsupervised Clustering

The resulting FPCA scores are used as input for unsupervised clustering using either the hdbscan (*dbscan*), fuzzy-clustering (*mfuzz)*, or kmeans (base R) algorithm. This procedure allowed data-driven identification of methylation trajectory subtypes without requiring prior knowledge of cluster number or shape.

### Simulated Data

A total of 3,000 synthetic CpG sites were simulated across 300 individuals, each assigned a random age between 1 and 100 years. Fifteen trajectory archetypes were implemented to represent a range of biologically plausible methylation patterns. These included: non-correlated, linear trajectories (increasing, decreasing), quadratic trends (increasing, decreasing), logarithmic transitions (increasing, decreasing), sigmoidal dynamics (increasing, decreasing) with inflection points at ages 25, 40, and 80, variance-increasing profiles, and non-monotonic patterns. For each of the 15 trajectory classes, 200 CpGs were simulated.

For all the functions except the variance-increasing, the age-specific methylation expectation (μ) was passed to a Beta distribution (via rbeta) to introduce variability while maintaining biological constraints (bounded between 0 and 1). For the variance-increasing function, age-dependent Gaussian noise was added directly to a mean of 0.5, with standard deviation increasing linearly from 0.01 (age < 25) to 0.3 (age = 100), mimicking stochastic methylation drift. Details are available in Supp. File.

### Benchmarking SNITCH

The benchmarking was done using the simulated patterns and their associated labels as ground truth. Fuzzy c-means clustering was performed using the *Mfuzz* package. The fuzzification parameter *m* was estimated using the function mestimate, and the optimal number of clusters was set to 11 after determining the minimum centroid distance across 3 repeated runs. Final cluster assignments were defined by the highest membership value. K-means was run with 25 restarts and centers = 10 by using the elbow method based on total within-cluster sum of squares (WCSS) and k.max = 20. HDBSCAN was applied using the *dbscan* package, the minimum cluster size was set to 5. The DICNAP pipeline was implemented as originally described^18^, except for the maximal number of clusters for K-means set to 20. SNITCH was run as previously described. The FPCA scores were subsequently used for unsupervised classification by the same three algorithms. The ‘*c’* parameter was set to 11 for fuzzy clustering, and centers = 10 for K-means. Each method’s cluster assignments were compared to ground-truth labels to compute ARI and AMI using the same functions from *aricode*. The results from the two best-performing methods were evaluated by a confusion matrix.

### Identification of sex-specific CpG Methylation Trajectories Using SNITCH

To evaluate aging-associated methylation trajectories independently of immune cell composition, we constructed three models differing in their treatment of cellular heterogeneity. In the Baseline (BL) model, we ran SNITCH with default parameters and without correcting for covariates. In the 7-cell and 12-cell models, we accounted for cell type proportions estimated using the *EpiDISH* package with the centDHSbloodDMC.m and cent12CT.m reference matrix, respectively, using the Robust Partial Correlations (RPC) method^24^. The 7 cell-matrix contained information on B-, NK-, CD4T, and CD8T-cells, Monocytes, Neutrophils, and Eosinophils. Building on the 7-cell reference matrix, the 12-cell model discriminated between naive and mature B-, CD4T-, and CD8T-cells, and added T-regulatory cells and Basophils. To identify groups of CpGs with similar aging dynamics, we applied both *Mfuzz* and *HDBSCAN* to the SNITCH-classified non-linear trajectories. Clustering results were assessed visually, and we retained only the *HDBSCAN* clusters for downstream analyses due to their superior trajectory homogeneity. Similarly, the optimal minPts parameter for HDBSCAN was determined by visual inspection of cluster resolution. The selected minPts values for the female BL, 7-cell, and 12-cell models were 5, 5, and 7, respectively. For the male models, they were 5, 8, and 4. Notably, HDBSCAN designates sparse or noisy data points as cluster 0. Upon reviewing the trajectories assigned to this group in the female 12-cell model, we observed two distinct patterns within cluster NL0. We therefore re-ran HDBSCAN on this subset to refine the classification, resulting in three clusters: NL10, NL11, and NL12. Then, for each NL cluster, we performed principal component analysis (PCA) and extracted PC1 to represent the dominant methylation pattern. A correlation matrix was then computed across all cluster PC1 using Spearman’s rank correlation. Clusters with correlations exceeding 0.90 were merged to reduce redundancy and capture shared underlying dynamics.

### Functional analysis

#### Classification of clocks and immune cells CpGs

CpGs were retrieved for the following clocks from the Biolearn resource^73^: Horvath v1, Hannum, PhenoAge, GrimAgeV2, DunedinPACE, YingCausAge, YingDamAge, YingAdaptAge, and Zhang_10. Only unique CpG identifiers were retained. We counted the occurrence of each clock CpGs in the different categories identified by SNITCH across our 3 models (BL, 7-cell, 12-cell) in both males and females. A similar analysis was performed with age-related CpGs (169 hypermethylated, 181 hypomethylated) shared across immune cells retrieved from Roy R. et al. ^33^.

#### Chromatin state enrichment

Chromatin states profiled in peripheral blood mononuclear cells (PBMCs) were obtained from the Roadmap Epigenomics Project ^34^and mapped to CpG probe IDs. Fisher’s exact tests were conducted to evaluate the enrichment of each chromatin state across aging trajectory classes against NC CpGs in both sexes. P-values were adjusted using the Benjamini-Hochberg method.

#### Pathway enrichment analysis

Both gsameth and gometh functions from the *missMethyl ^43^*R package were used to perform Over Representation Analysis (ORA). These functions account for the differing numbers of CpG probes per gene on the Illumina EPIC array, thereby reducing bias inherent to standard enrichment tools when applied to DNAm data. The gsameth function allows testing a list of CpGs against a custom gene set. We retrieved 2 custom gene sets from the Molecular Signatures Database (MSigDB v2024.1/v2025.1): Human Phenotype Ontology^85^ (HPO - <u>c5.hpo.v2024.1.Hs</u>.entrez.gmt and Reactome Pathways ^86^(c2.cp.reactome.v2025.1.Hs.entrez.gmt). The Gene Ontology^87^ and KEGG ^88^databases were tested with the gometh function. We used the set of CpG sites corresponding to each DNA methylation cluster, excluding the non-correlated “NC” sites, as the test set, and the full set of tested CpG sites across all clusters served as the background. Analyses were conducted independently for male and female clusters. The analysis was replicated for the CpGs of each peak identified during the DEswan analysis.

#### Motif enrichment analysis

The motif enrichment tool from EWAS datahub ^89^ was used to perform motif enrichment analysis on each cluster identified in females and males. This tool uses a centered 500bp window on the CpG of interest to perform Hypergeometric Optimization of Motif EnRichment (HOMER)^90^. Motifs with a qvalue < 0.05 were considered significant. The analysis was replicated for the CpGs of each peak identified during the DEswan analysis.

#### Trait association - Cancer

The cohort used to assess the sex-specific biomarker potential of cluster eigenvalues has been described elsewhere. Briefly, the EPIC-Italy^59^ study includes DNA methylation data collected at baseline and linked to up to 14 years of prospective follow-up. Cancer cases were annotated with time to diagnosis and cancer type, enabling time-to-event analyses. We performed sex-stratified analyses using DNAm of patients preprocessed as described above. Individuals without a cancer event were treated as right-censored. For these, the time to diagnosis was imputed using the maximum observed follow-up time among cases. For each patient, we computed their NL cluster eigenvalues as described above.

Importantly, this step was restricted to the CpGs common to both the EPIC and 450K arrays, resulting in smaller sets of CpGs per cluster (Supp.Table). Immune cell-type proportions were estimated from whole blood methylation profiles using the EpiDISH ^24^algorithm (RPC method) with a 12-cell reference panel. The coxph function from the *survival* package was used to fit cox proportional hazards models using “Surv(time_to_diagnosis, status)” as the outcome, with eigenvalues, age, and immune cell proportions as predictors. Logistic regression models (glm(family=binary)) using cancer status as the binary outcome were also fit for comparison. Model coefficients were exponentiated to yield hazard ratios (HR) and odds ratios (OR), with 95% confidence intervals. To assess survival differences, samples were stratified into tertiles based on the significant eigenvalues (Low, Mid, High). Kaplan– Meier curves were generated using the survfit function and plotted with ggsurvplot() (log-rank p-value and 95% CI shown) from the *survminer* package. Significance was considered as FDR < 0.05.

#### Trait association - Inflammation

Separately in males and females, we computed the “eigenvalue” of each cluster, corresponding to that cluster’s first principal component (PC1), by performing principal component analysis (PCA) on centered and scaled methylation beta values across CpGs within that cluster. For each sample, we used the *ComputeCRPscore*^62^ function based on a predefined set of weighted CpGs to generate an estimation of CRP protein levels as a surrogate for inflammation^91^. To test the independent contribution of age-associated methylation modules to CRP variation, we constructed a series of nested linear models:

- **Model 1** included **age** as the only predictor; CRP ∼ age
- **Model 2** included age and the eigenvalue of the NC (non-correlated) CpGs; CRP ∼ age + NC
- **Model 3 (full model)** included age, the NC eigenvalue, and eigenvalues of the VI, LI, LD, and NL clusters. CRP ∼ age + NC + VI + LI + LD + NLi + … + NLn

We compared models using ANOVA to evaluate whether the addition of SNITCH-derived modules significantly improved the explanation of CRP variability beyond age and (non-correlated) methylation patterns. Model fit and the significance of individual predictors were assessed using standard linear modeling statistics and p-values < 0.05.

#### Wave of aging – DEswan analysis

We performed a DEswan analysis as previously described^1,6^. DEswan works as a sliding window, where at each specified time point centered at the middle of the window, CpGs at both ends are compared using a Wilcoxon test for differential methylation. We restricted analyses to age-associated CpGs previously identified (excluding NC CpGs). We used windows centered from 31 to 78 years in 2-year steps with a 15-year bucket size. To account for cell composition, we estimated per-sample blood cell-type proportions with EpiDISH using a 12-cell reference (cent12CT) and included these estimates as covariates in all models (as described above). P-values were adjusted for multiple testing by the Benjamini-Hochberg method, and significance was considered at FRD < 0.05. To ensure the robustness of the findings, we only considered peaks that were conserved at FDR < 0.01 and 0.001 (Figure 4B). We adapted the initial R script to allow parallelization across CpGs using parLapply.

## Supplementary Figures

**Supp. Figure 1:**
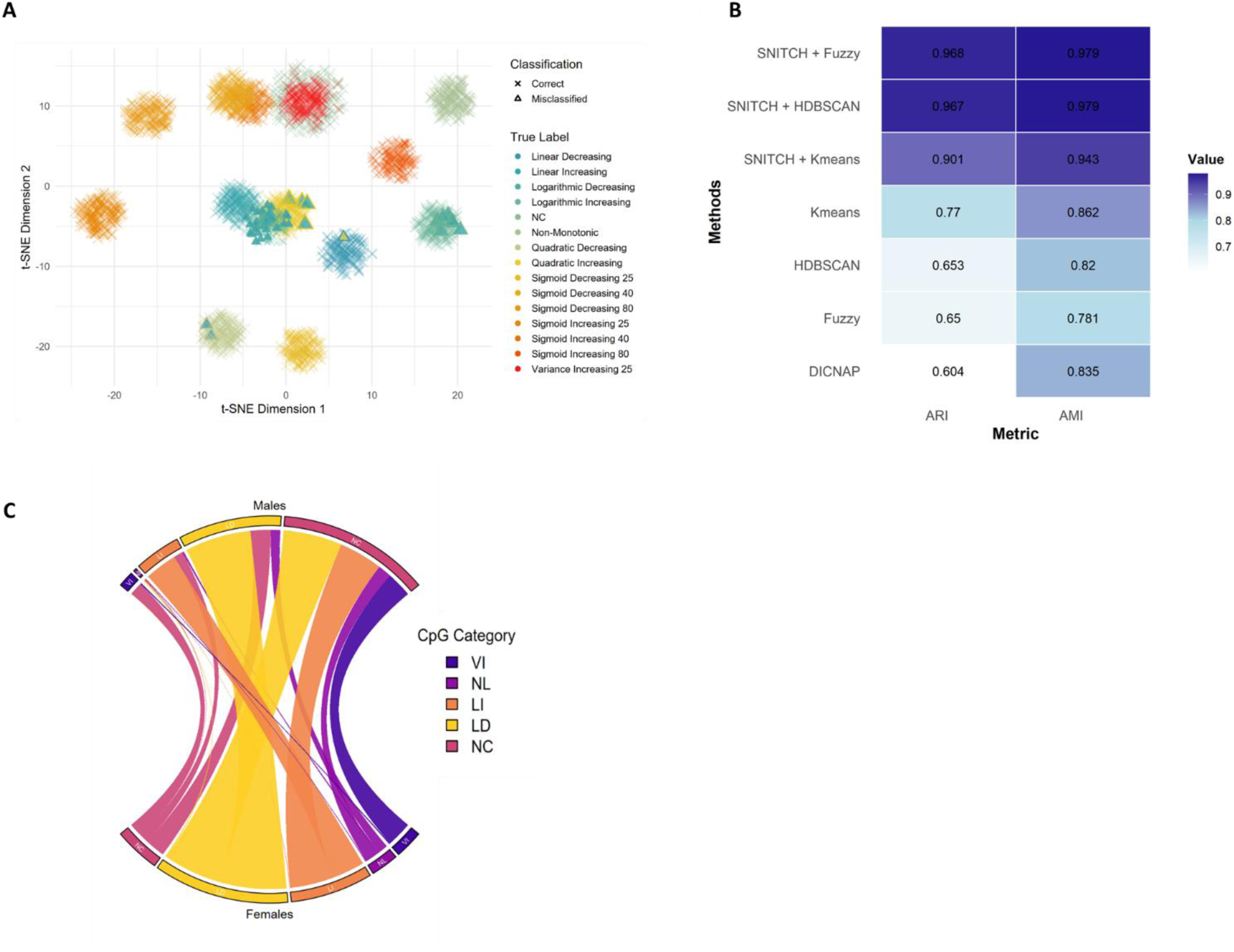
**A**, T-SNE representation of the simulated CpGs to illustrate their true and assigned labels after SNITCH. **B**, Benchmark of SNITCH compared to stand-alone unsupervised clustering methods and DICNAP. **C**, Conserved CpGs between male and female clusters.

**Supp. Figure 2:**
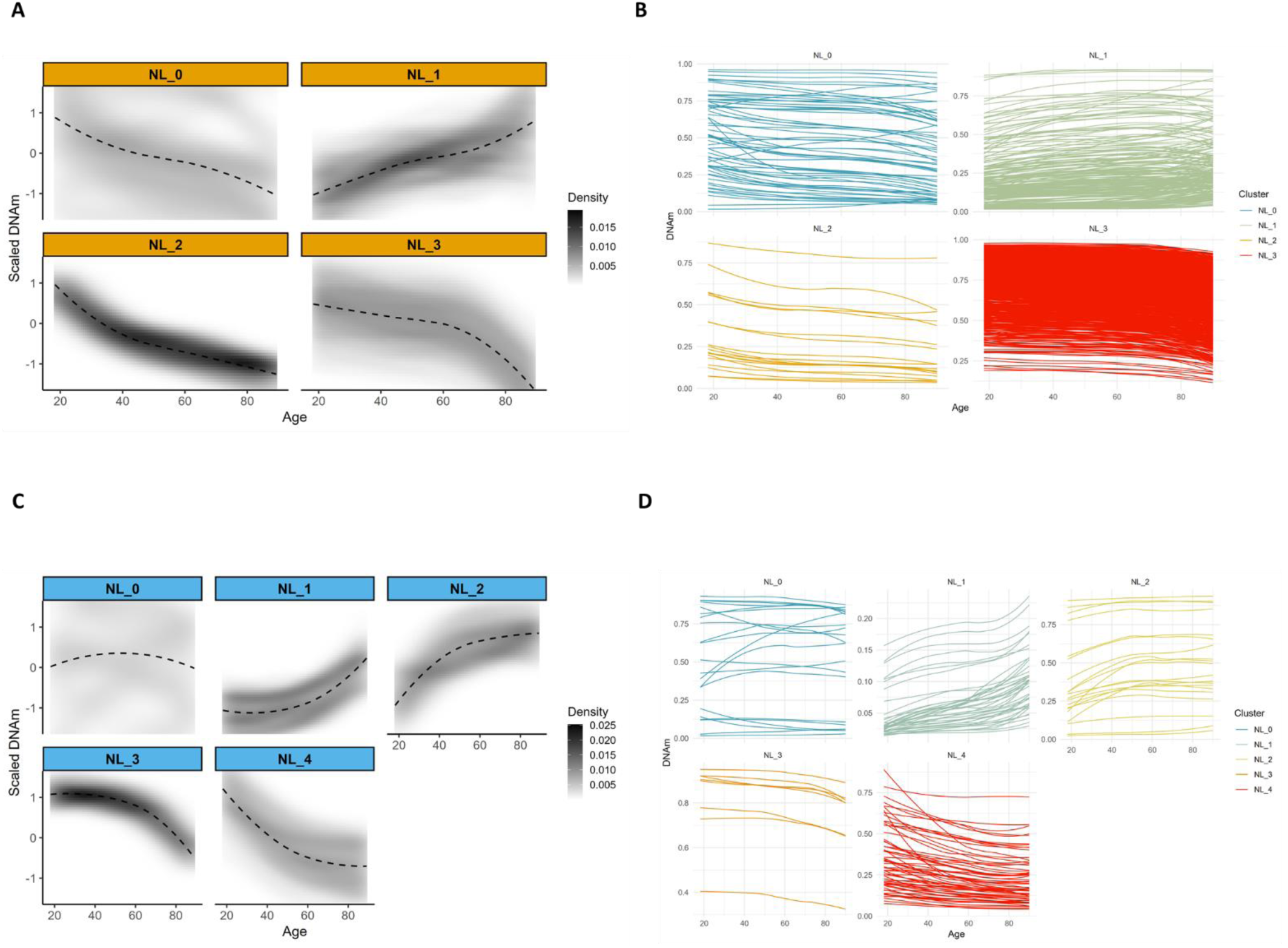
**A**, Initial non-linear clusters identified in females. Beta values were centered and scaled prior to FPCA and unsupervised clustering and are used to better illustrate the patterns. **B**, Unscaled beta values in the initial females’ non-linear clusters.: **C**, Initial non-linear clusters identified in males. Beta values were centered and scaled prior to FPCA and unsupervised clustering and are used to better illustrate the patterns. **B**, Unscaled beta values in the males’ initial non-linear clusters.

**Supp. Figure 3:**
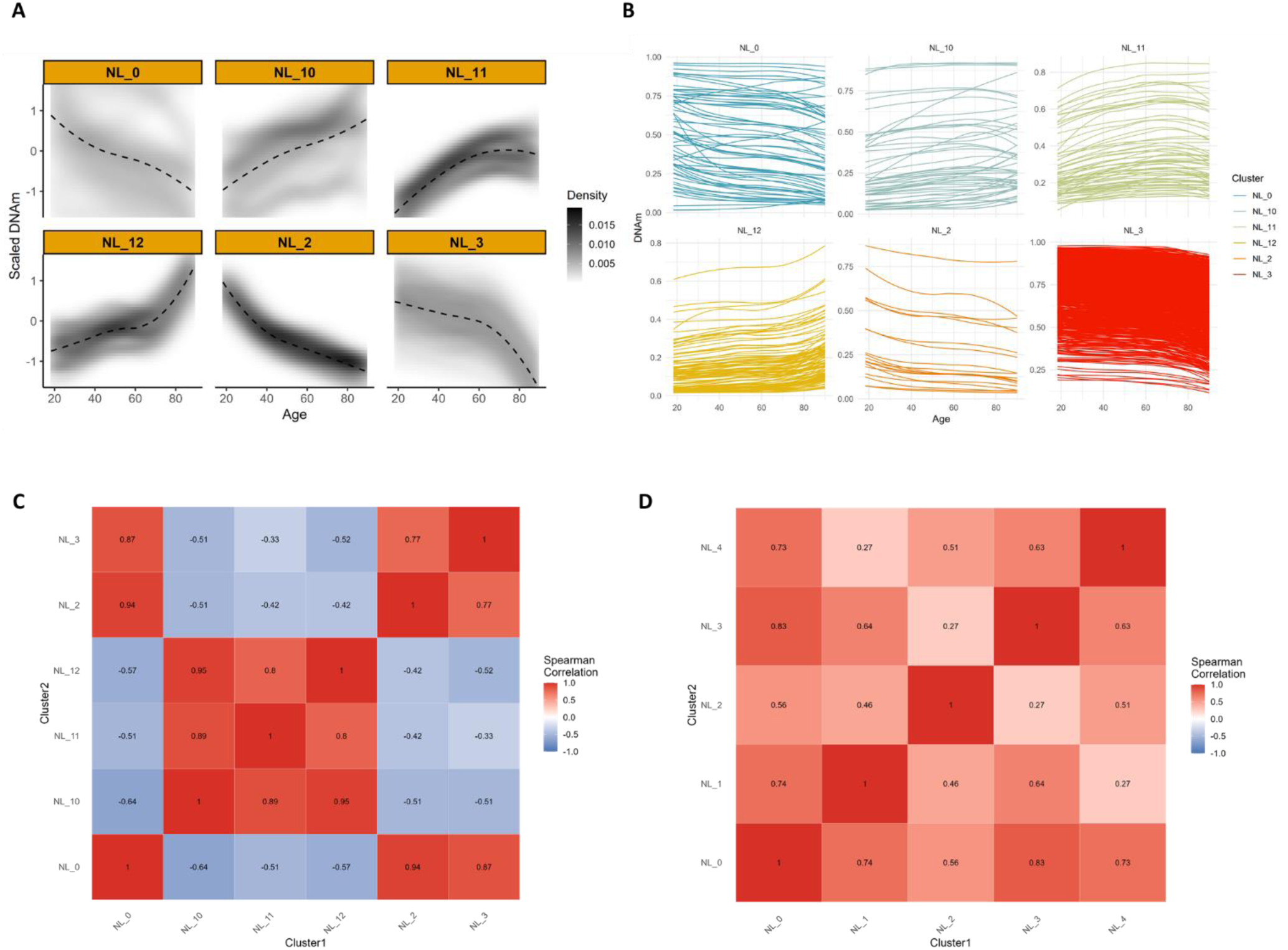
**A**, Non-linear clusters identified in females after the reclassification of NL1. Beta values were centered and scaled prior FPCA and unsupervised clustering and are used to better illustrate the patterns. **B**, Unscaled beta values in the females’ non-linear clusters after the reclassification of NL1. **C**, Correlation matrix between the NL clusters eigenvalues in females. **D**, Correlation matrix between the NL clusters eigenvalues in males.

**Supp. Figure 4:**
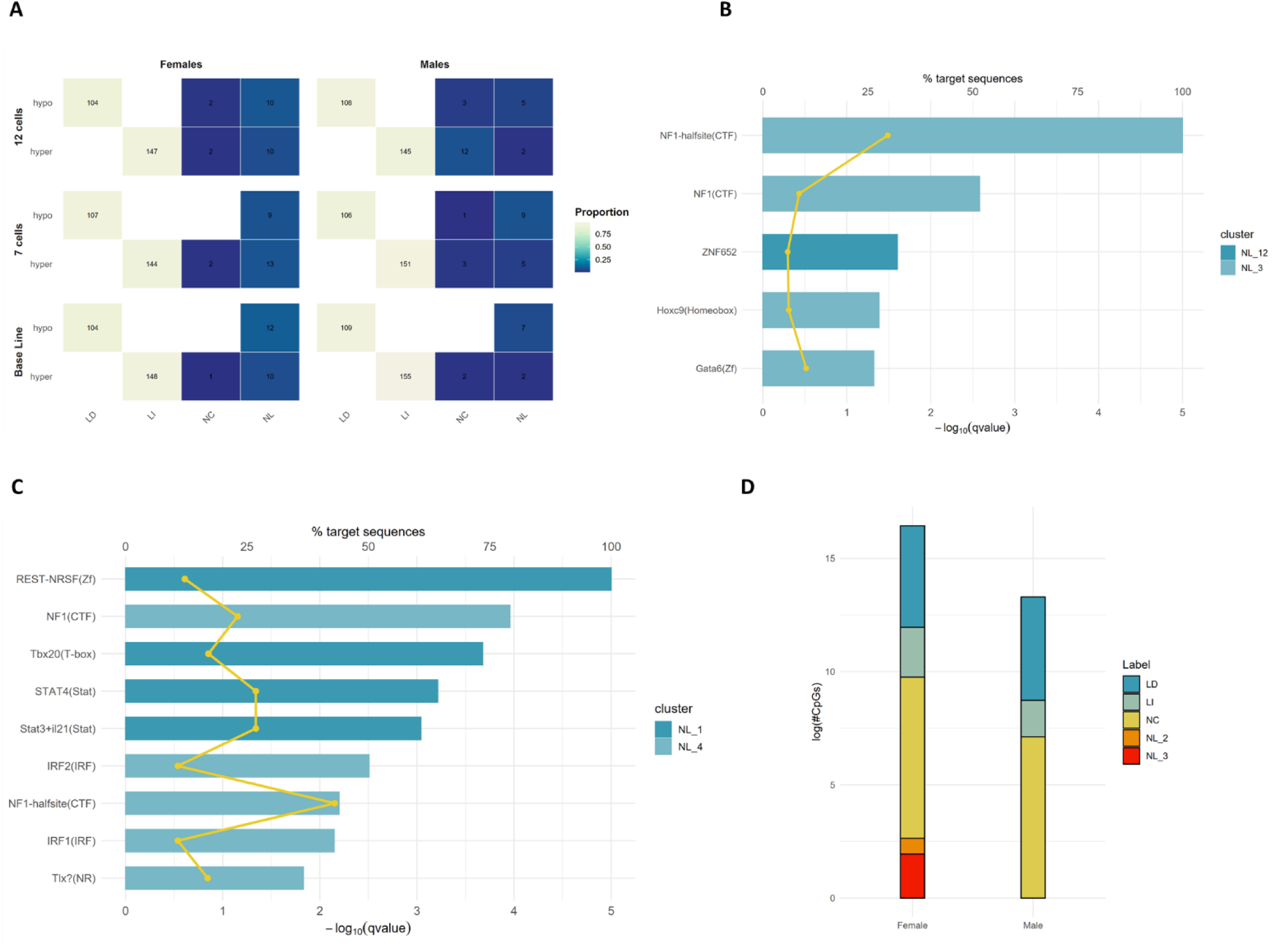
**A**, Enrichment of CpG labels across age-CpGs identified in the EWAS study by Roy R. et al.. **B**, Cluster-wise motif enrichment analysis among females NL CpGs. **C**, Cluster-wise motif enrichment analysis among females NL CpGs. **D**, Distribution of the CpGs used to estimate C-Reactive Protein levels among males and females aging classification.

**Supp. Figure 5:**
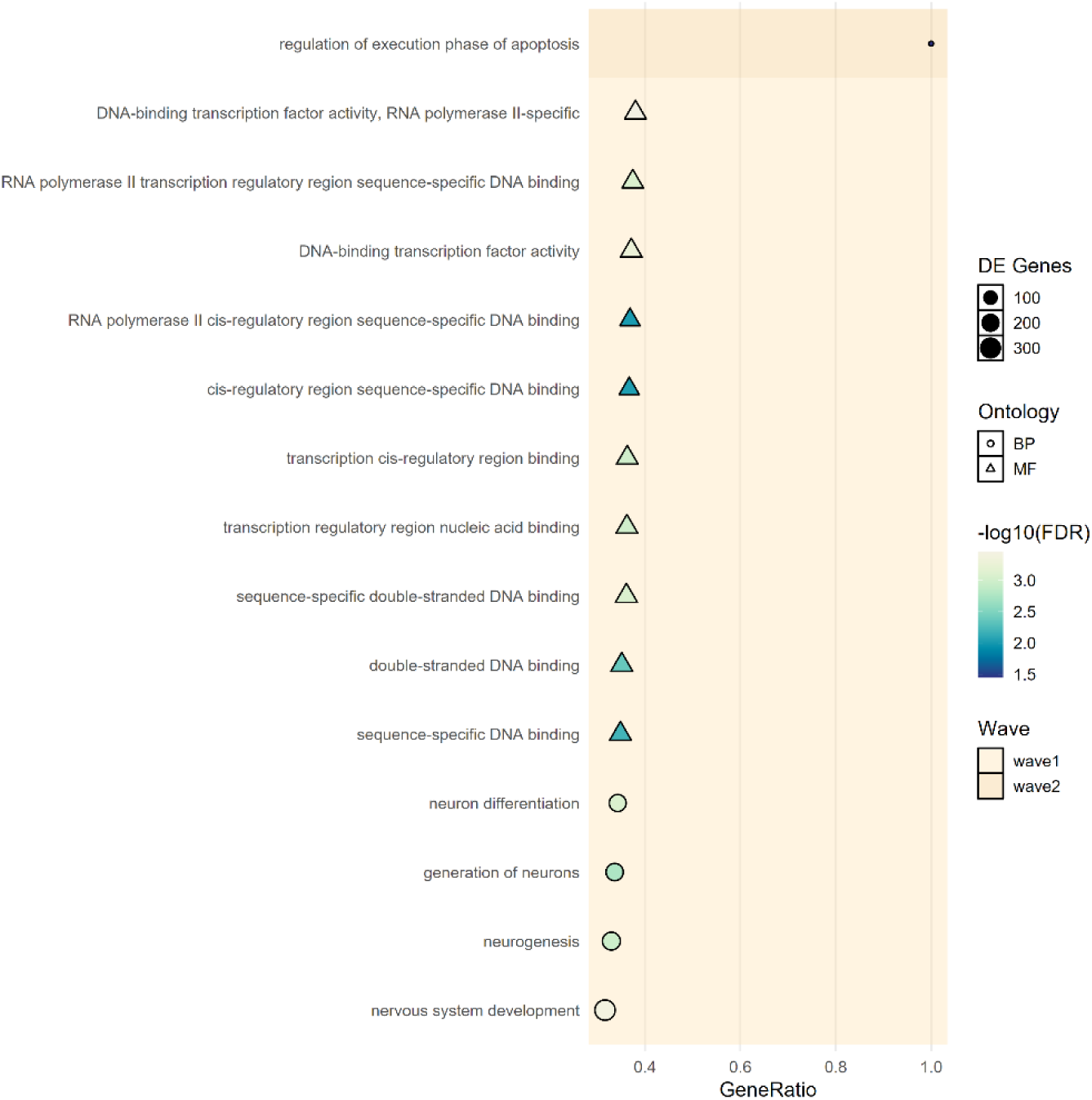
Pathway enrichment analysis performed among common CpGs identified across waves of dysregulation in females.

